# Machine Learning Framework for Fully Automatic Quality Checking of Rigid and Affine Registrations in Big Data Brain MRI

**DOI:** 10.1101/2020.10.23.352781

**Authors:** Sudhakar Tummala, Niels K. Focke

## Abstract

Rigid and affine registrations to a common template are the essential steps during pre-processing of brain structural magnetic resonance imaging (MRI) data. Manual quality check (QC) of these registrations is quite tedious if the data contains several thousands of images. Therefore, we propose a machine learning (ML) framework for fully automatic QC of these registrations via local computation of the similarity functions such as normalized cross-correlation, normalized mutual-information, and correlation ratio, and using these as features for training of different ML classifiers. To facilitate supervised learning, misaligned images are generated. A structural MRI dataset consisting of 215 subjects from autism brain imaging data exchange is used for 5-fold cross-validation and testing. Few classifiers such as *kNN*, *AdaBoost*, and *random forest* reached testing F1-scores of 0.98 for QC of both rigid and affine registrations. These tested ML models could be deployed for practical use.

## 1. INTRODUCTION

Given the increasing availability of publicly accessible repositories for brain magnetic resonance imaging (MRI) data, researchers have been enabled to process larger number of subjects in neuro-scientific applications; such studies are often phrased “big data” MRI studies [1, 2]. Some of these publicly available brain MRI big data sets include HCP (aging, young adult, development), IXI, ADNI, ABIDE, OASIS, UK biobank, and NAKO Germany [3]. Due to advances in high-performance computing and the availability of these big data, brain studies on a larger population are now getting feasible [4, 5]. Structural MRI is the prime modality for the studies involving voxel-based morphometry (VBM) of the whole brain, and surface-based morphometry of cortical gray matter structures [6–8].

Quality control (QC) has become an important step starting from assessing raw image quality until the final post-processing. Checking the data for quality before pre-processing is the first stage of QC and there have been several studies proposing different metrics for quantitative assessment of raw image quality [9, 10].

After initial QC has passed, these acquired raw structural MRI images have to undergo several standard pre-processing steps such as reorientation to a standard template, cropping, bias correction, and alignment of the images in the subject space to a common space. Transforming images to a common space is one of the crucial steps in the pre-processing stage to make them better suitable for further processing such as VBM. These transformations could involve both rigid and affine registrations with 6 and 12 degrees of freedom respectively. QC of these registrations is necessary to make sure that they are suitable for subsequent analysis. In studies involving data in a few hundred, manual checking is the common solution. However, in big data studies that involve tens of thousands of images, manual inspection will be a time-consuming process and moreover the manual process could be prone to inter/intra-observer rating errors especially with doubtful cases. Hence, this raises the need for a fully automatic quality control mechanism which potentially reduces the manual intervention to the possible minimum.

Machine learning (ML) has been in wide use in medical imaging for several tasks such as assessment of image quality, brain mapping, and disease diagnosis and prognosis [11–13]. Coming to registration quality, the traditional similarity cost functions primarily in use are the sum of squared differences, normalized cross-correlation (*NCC*), joint entropy-based methods such as normalized mutual information (*NMI*) and correlation ratio (*CR*) [14]. However, computing them locally and combining them by employing ML classifiers may result in better accuracy compared with accuracy using the individual cost functions alone.

To our knowledge, this is the first study aiming at the development of different supervised ML models that could be deployed for use in big data structural MRI pre-processing.

## 2. METHODS

This section describes the dataset used for pre-processing, generation of misaligned images for supervised learning, and computation of local cost values, cross-validation, and testing of different ML classifier models.

### 2.1. Dataset

The data for cross-validation and testing consisted of a subset of high-resolution T1-weighted MRI images of 215 subjects extracted from publicly available autism brain imaging data exchange (ABIDE I) [2]. Out of 215 subjects, 110 subjects are with autism. These images were acquired using a 3.0 T Siemens scanner with magnetization prepared rapid acquisition gradient echo sequence in the sagittal plane.

### 2.2. Pre-processing

The data is pre-processed under the Nipype environment by calling the required third-party interfaces such as ANTS, FSL, Freesurfer, and other required software [15]. Firstly, the raw images are reoriented to standard space and cropped to remove the neck region using the FSL tools *fslreorient2std*, *robustfov* respectively [16]. Further, the cropped images are bias-corrected to remove the low-frequency intensity variations due to the inhomogeneity of the scanner magnetic field using the N4 method from ANTS [17]. Lastly, the images are aligned to the 1×1×1 mm^3^ standard T1-weighted FSL template both rigidly and affinely using the FSL *flirt* interface. After rigid and affine registration, the images are manually checked to make sure that they are correctly aligned to the template.

### 2.2. Generation of misaligned images

Since a supervised learning approach is used in training ML models, misaligned rigid and affine test images are created by decomposing the corresponding transformation matrix into scales, translations, and rotations; changing and applying these modified matrices to align the corresponding images in subject space to the common template. Ten test images (five each for rigid and affine) are generated for each correctly aligned image by altering the matrix parameters in different combinations to have a variety of misaligned images. Fig. 1 showing the generated rigid misaligned images for a subject.

**Fig. 1.**
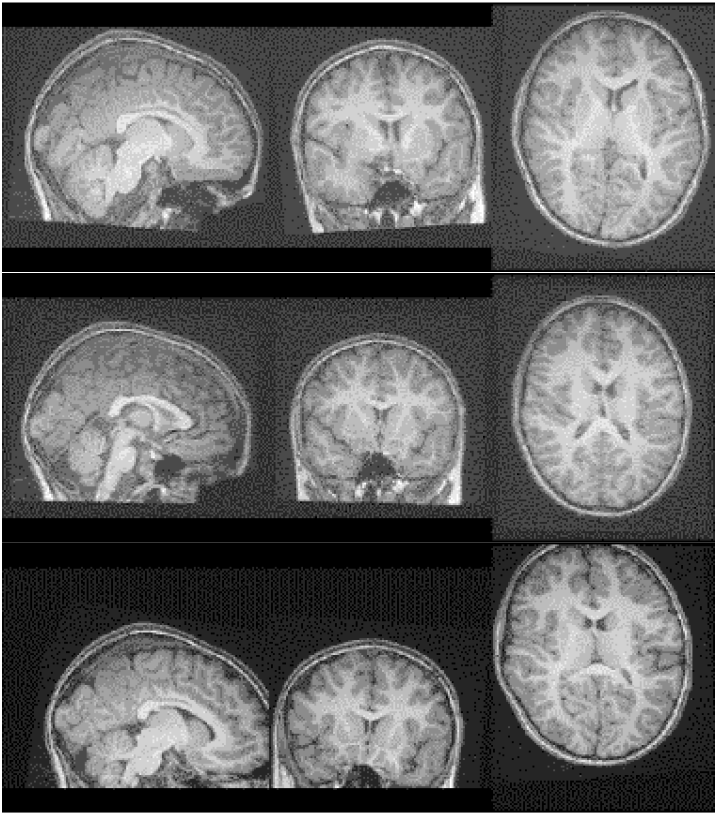
Generated rigid misaligned images shown in slices in the top (little variation) and bottom (more variation) rows, middle row is showing the correctly aligned image slices for comparison.

### 2.3. Computation of local cost values

Using a custom-written python (version 3) script via anaconda, the cost values that indicate the quality of registration between the moving image and the template are computed locally using the similarity functions such as *NCC*, *NMI*, and *CR* by defining a 3×3×3 volume of interest (VOI). The quantifications are restricted to the brain region by using the brain mask of the standard T1-weighted FSL template. The final cost value (*F_C_*) is the average of the local cost values obtained by moving the VOI with a stride of 3 and computing the cost values at each stride as described by the equation (1).

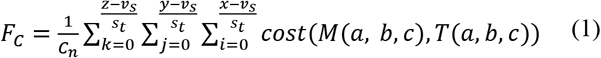

In equation (1), *a* = *i* * *s_t_*: ((*i* * *s_t_*) + *v_s_*), b = *j* * *s_t_*: ((*j* * *s_t_*) + *v_s_*), and *c* = *k* * *s_t_*: ((*k* * *s_t_*) + *v_s_*) indicate indices ranges of the moving image *M* and template *T*. *C_n_* is the total number of local cost computations, *v_s_* is the size of VOI and *s_t_* is the stride, and the *cost* is anyone of the costs given in equations (2), (3), and (4) below. Lastly, *x, y,* and *z* represent the sizes of the template (or moving) in the three directions. Although the size of VOI and stride are free parameters, they are restricted to 3 for the purpose of this study. The local cost *NCC* is computed as follows:

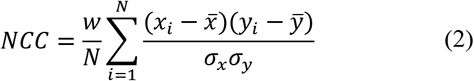

Above, *N* is the total number of non-zero voxels in either moving VOI or template VOI, *x_i_* is the i^th^ intensity and 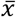 is the mean intensity of the moving VOI and similarly, *y_i_* is the i^th^ intensity and 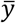 is the mean intensity of the template VOI, σ_*x*_ and σ_*y*_ are standard deviations of intensities of moving and template VOIs respectively. Further, the local cost *NMI* is computed using the expression in equation (3).

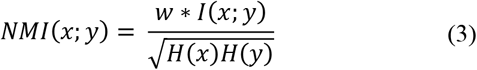

Where *I*(*x;y*) = *H*(*x*) + *H*(*y*) − *H*(*xy*) is the mutual information between *x* (moving VOI) and *y* (template VOI), and *H*(*x*) and *H*(*y*) are entropy values of VOIs *x* and *y* respectively. *H*(*xy*) is the joint entropy of *x* and *y*. Finally, the local cost *CR* is computed using the following expression:

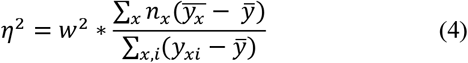

In equation (4), *y*_*xi*_ is the *i*^th^ intensity value in category *x* (category is either moving VOI or template VOI) *n*_*x*_ is the number of intensities in category *x*, 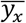 is the average of category *x* and 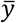 is the mean of intensities of both moving and template VOIs together. Finally, the *CR* is just *η*.

In equations (2), (3), and (4), *w* is the weighting factor which is computed using equation (5) to reduce the say of the cost values computed at boundaries of the brain on the final cost value *F_C_* since the VOIs at the border could contain zero intensities. In eq. (5), *c* is a constant that is fixed to 1000 and *g* is the number of non-zero intensities in the respective VOI.

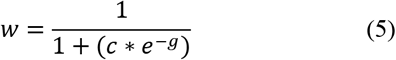

### Training of ML classifiers

Python based Scikit-learn (*sklearn*) module is used for fitting/training of ML classifier models [18]. Before feeding to ML classifiers, the three cost values *NCC*, *NMI* and *CR* are rescaled to have values between zero to one. The ML classifiers that are cross-validated and tested include linear discriminant analysis (*LDA*), Gaussian naïve Bayes (*GNB*), linear support vector classifier (*SVC*), k-nearest neighbors (*kNN*, *k* is chosen as 15), random forest (*RF*, 100 decision trees), and Adaptive Boosting (*AdaBoost* with 100 decision stumps). The data is divided into two groups where one group is used for repeated (100 repetitions) stratified 5-fold cross-validation (*CV*) and the other group is utilized for testing (Table 1). The area under the ROC curve (AUC) and F1-scores are calculated using the *sklearn* module to indicate the performance of each classifier during *CV* as well as testing.

**Table. 1.** Number of images used for *CV* and testing. N_1_ includes both healthy and diseased, N_2_ is the number of diseased.

| | *CV* $N_1$ ($N_2$) | Testing $N_1$ ($N_2$) |
| --- | --- | --- |
| **Correctly aligned** | 170 (87) | 45 (23) |
| **Misaligned** | 850 (435) | 225 (115) |

## 3. RESULTS

The F1-scores during *CV* and testing for identifying the misaligned registrations using different ML classifiers is given in Table 2 for both rigid and affine. The cross-validated ML models achieved the testing F1-scores in the range of 0.914–0.988 and the AUC values between 0.972–0.991 for identifying misaligned rigid registrations. Fig. 2 shows the true positive rate (sensitivity) and false positive rate (1-specificity) ROC curves for different classifiers that are validated on the test set for finding faulty rigid registrations.

**Table. 2.**
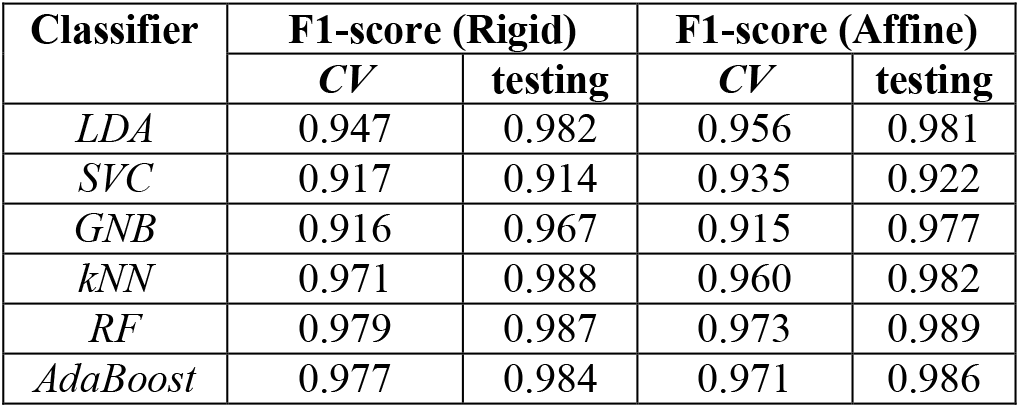
F1-scores during *CV* and testing of different ML classifiers for checking both rigid and affine registrations.

| Classifier | F1-score (Rigid) |  | F1-score (Affine) |  |
| --- | --- | --- | --- | --- |
|  | *CV* | testing | *CV* | testing |
| *LDA* | 0.947 | 0.982 | 0.956 | 0.981 |
| *SVC* | 0.917 | 0.914 | 0.935 | 0.922 |
| *GNB* | 0.916 | 0.967 | 0.915 | 0.977 |
| *kNN* | 0.971 | 0.988 | 0.960 | 0.982 |
| *RF* | 0.979 | 0.987 | 0.973 | 0.989 |
| *AdaBoost* | 0.977 | 0.984 | 0.971 | 0.986 |

**Fig. 2.**
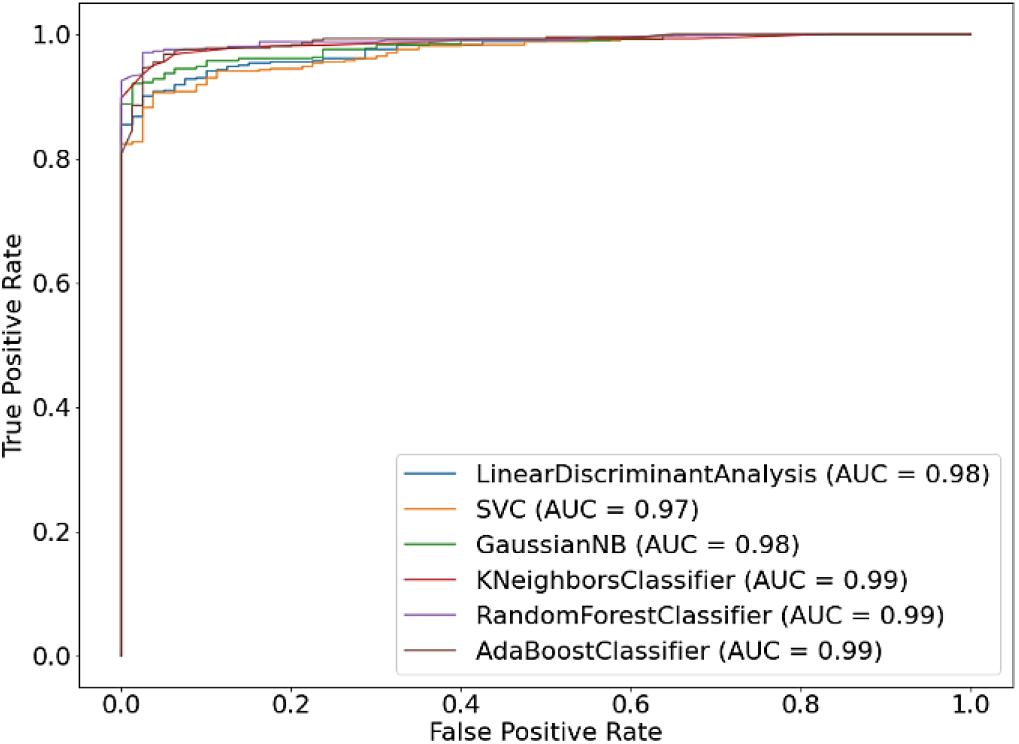
Graph of true positive rate vs false positive rate for different ML classifiers tested for rigid registrations.

Similarly, for affine, the cross-validated ML models reached the testing F1-scores and AUC values in the ranges of 0.922–0.989 and 0.989–0.993 respectively. Fig. 3 shows the ROC curves for different classifiers that are validated on the test set for finding incorrect affine registrations.

**Fig. 3.**
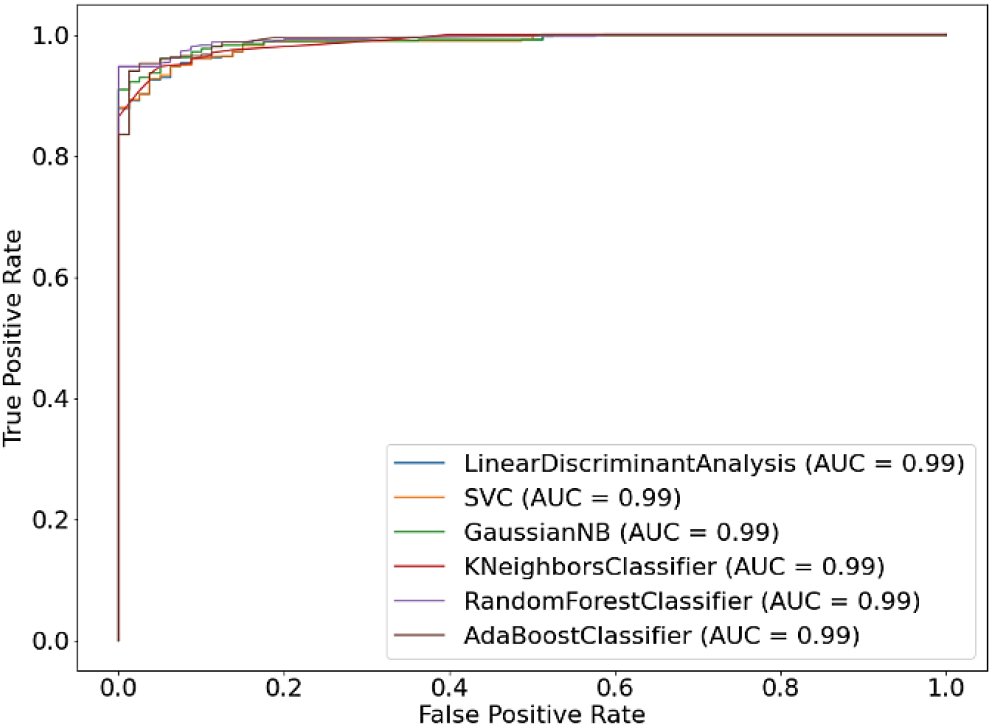
Graph of true positive rate vs false positive rate for different ML classifiers tested for affine registrations.

## 4. DISCUSSION

Here, we developed a fully automatic framework using different ML classifier models for QC of rigid and affine registrations during structural MRI pre-processing. The classifier selection could be based on F1-score as the number of images in the two classes is heavily imbalanced during *CV* an testing. From Table 2, figs. 2 and 3, it is evident that all classifiers performed well, overall *kNN*, *RF*, and *AdaBoost* classifiers demonstrated both better testing F1-scores and AUC values. Another aspect to consider here is that checking the quality of alignment by computing the cost values locally using a moving VOI might have led to the anticipated F1-sores and AUC values on the test set. Since a stride of 3 is applied, which significantly reduced the computational time to around a minute for each local cost value, thus making it computationally efficient. The tested ML models could be deployed using the *pickle* module for practical use. The whole processing pipeline is available at <u>tummala-github</u>.

## 5. CONCLUSION

The developed ML models could be deployed for fully automatic quality checking of rigid and affine registrations in big data brain structural MRI pre-processing. Since the data is validated both on healthy and diseased brains, the trained models may be capable of identifying the misalignments both in health and disease. Future work could involve testing and improving these developed classifier models by using different larger cohorts and may be extended to deal with non-linear registrations as well. Further, the development of deep neural nets could be considered to eliminate the need for computation of local cost values explicitly. Also, a single framework could be developed to deal with all types of structural images such as T1-weighted, T2-weighted, FLAIR, and proton density.

## 6. COMPLIANCE WITH ETHICAL STANDARDS

This research study was conducted retrospectively using human subject data made available in open access by (ABIDE I). Ethical approval was not required as confirmed by the license attached with the open access data.

## 7. ACKNOWLEDGMENTS

- This work was supported by research funds from the Department of Science, state of Lower Saxony, Germany.
- The authors have no relevant financial or non-financial interests to disclose.

